# Fear conditioning prompts sparser representations of conditioned threat in primary visual cortex

**DOI:** 10.1101/2020.01.16.909085

**Authors:** Siyang Yin, Ke Bo, Yuelu Liu, Nina Thigpen, Andreas Keil, Mingzhou Ding

**Affiliations:** J. Crayton Pruitt Family Department of Biomedical Engineering, University of Florida, Gainesville, FL 32611; Center for Mind and Brain, University of California, Davis, CA 95618; Department of Psychology, University of Florida, Gainesville, FL 32611

**Keywords:** fear conditioning, alpha oscillations, visual representation, sparsification, attention

## Abstract

Repeated exposure to threatening stimuli alters sensory responses. We investigated the underlying neural mechanism by recording simultaneous EEG-fMRI from human participants viewing oriented gratings during Pavlovian fear conditioning. In acquisition, one grating (the CS+) was paired with a noxious noise, the unconditioned stimulus (US). The other grating (CS-) was never paired with US. In habituation, which preceded acquisition, and in final extinction, the same two gratings were presented without the US. Using fMRI-BOLD multivoxel patterns in primary visual cortex during habituation as reference, we found that during acquisition, aversive learning selectively prompted systematic changes in multivoxel patterns evoked by the CS+. Specifically, CS+ evoked voxel patterns in V1 became sparser as aversive learning progressed, and the sparse pattern was preserved in extinction. Concomitant with the voxel pattern changes, occipital alpha oscillations were increasingly more desynchronized during CS+ (but not CS-) trials. Across acquisition trials, the rate of change in CS+-related alpha desynchronization was correlated with the rate of change in multivoxel pattern representations of the CS+. Furthermore, alpha oscillations co-varied with BOLD in the right temporal-parietal junction, but not with BOLD in the amygdala. Thus, fear conditioning prompts persistent sparsification of threat cue representations, likely mediated by attention-related mechanisms.

## Introduction

Accurate detection and evaluation of threat and danger is crucial to survival. The mammalian brain has evolved mechanisms that bias perceptual systems towards sensory cues that predict aversive outcomes (Pessoa and Adolphs, 2010). For example, neurons in human primary visual cortex (V1) alter their tuning behavior to selectively amplify visual threat cues (Miskovic and Keil, 2012). Across species, sensory neurons in rodents also undergo selective plasticity to better represent threat cues, both in the visual cortex (Shuler & Bear, 2006) and in the auditory cortex (Weinberger, 2004). These observations suggest that associative learning of contingencies between a conditioned visual stimulus (CS+) and an aversive unconditioned stimulus (US) prompts changes in the neural representation of CS+.

Electrophysiological studies in humans have demonstrated selective amplification of primary visuocortical responses to the CS+ compared to control stimuli never paired with US (CS-; Thigpen et al., 2017). Paralleling conditioned auditory receptive field plasticity in rats (Headley and Weinberger, 2011), these sensory changes can be characterized as selectively heightened population gain for the critical CS+ feature (McTeague et al., 2015). What is not known, however, is how the visuocortical representation of a threat cue is changing as learning progresses. Possible hypotheses include (1) an increase in visuocortical population gain when viewing a threat-associated cue (Morris et al., 1998; Phelps et al., 2006), and (2) the emergence of efficient, sparse, highly connected visuocortical networks, through Hebbian mechanisms (Simoncelli and Olshausen, 2001; Miskovic and Keil, 2012; Headley and Weinberger 2013). Testing of these competing views has been elusive. In terms of fMRI, sparsification would be reflected in increasingly differential representations only for CS+, not for CS-, during fear conditioning, with decreasing numbers of voxels contributing to the representation of the CS+. By contrast, heightened population responses when viewing the CS+ would result in heightened BOLD in a larger number of voxels. We addressed these competing notions by measuring fMRI and quantifying the evolutions of fMRI patterns evoked by conditioned stimuli.

Autonomic orienting responses (e.g., heart rate) to conditioned threat attenuates along with CS+-related response in the limbic structures (Yin et al., 2018). To what extent this process is accompanied by enhanced attentional orienting is not clear. We measured EEG concurrently with BOLD so that EEG alpha band activity (8-12 Hz) could be used as an index of visual attention engagement. Transient suppression of spectral power in the alpha band (i.e., event-related desynchronization or ERD) has been taken to index attentive engagement of visual cortex in processing task-relevant stimuli (Klimesch, 2012; Zumer et al., 2014), and the more task-relevant the stimuli the stronger the alpha ERD (Klimesch et al., 2011; Auksztulewicz et al., 2107). We hypothesized that as threat cues acquire increased task-relevance through conditioning, alpha power would show greater ERD after CS+ stimuli compared to CS- stimuli, and this effect would become stronger as learning progressed.

BOLD responses in V1 and alpha ERD are both modulated by attention control networks (Posner and Gilbert, 1999; van Diepen et al., 2016), and alpha power reductions index target selection during a range of selective attention tasks (e.g., Rohenkohl and Nobre, 2011). However, in fear conditioning, the structures contributing to the selective visuocortical changes remain unclear. Two potential sources of modulatory bias signals are the ventral attention network (VAN), and limbic emotion-modulated circuits centered around the amygdala (McHugo et al., 2013; Yates et al., 2010). The VAN, including right ventrolateral prefrontal cortex (vlPFC) and right temporal-parietal junction (rTPJ), is involved in directing attention toward salient stimuli (Corbetta et al., 2002). The amygdala encodes information about the motivational significance of visual stimuli (Amaral et al., 2003; Paton et al., 2006) and may modulate visual cortex through connections with the basal forebrain (Peck et al., 2014) or with parietal and temporal cortex (Amaral et al., 2003; Keil et al., 2009). We examined these competing possibilities by correlating alpha power fluctuations with fMRI from rTPJ and amygdala.

## Materials and Methods

### Experimental Procedure

#### Participants

The experimental protocol was approved by the Institutional Review Board (IRB) of the University of Florida. Eighteen healthy college students (aged 17–33 years, nine females) provided written informed consent and participated in the study. The participants were either paid or given course credits in accordance with IRB guidelines.

#### Stimuli

Two Gabor patches (sine wave gratings filtered with a Gaussian envelope, Michelson contrast = 1) with the same spatial frequency (1.5 cycles/degree), differing only in orientation (45° and 135°), were designated as CS+ and CS-respectively; they were not counterbalanced across subjects. See Figure 1. Both stimuli were projected onto a back-illuminated screen (60 cm X 60 cm) placed 230 cm away from the participant’s head and viewed through a set of prismatic glasses attached to the radio frequency head coil. The US was a 1-second human scream delivered by a MRI compatible headphone at around 95dB. For CS-trials and CS+ trials where CS+ and US were not paired, the Gabor patches were shown for 1 second. For CS+ trials where CS+ and US were paired, the US started 0.5 second following CS+ onset and co-terminated 1 second later.

**Figure 1:**
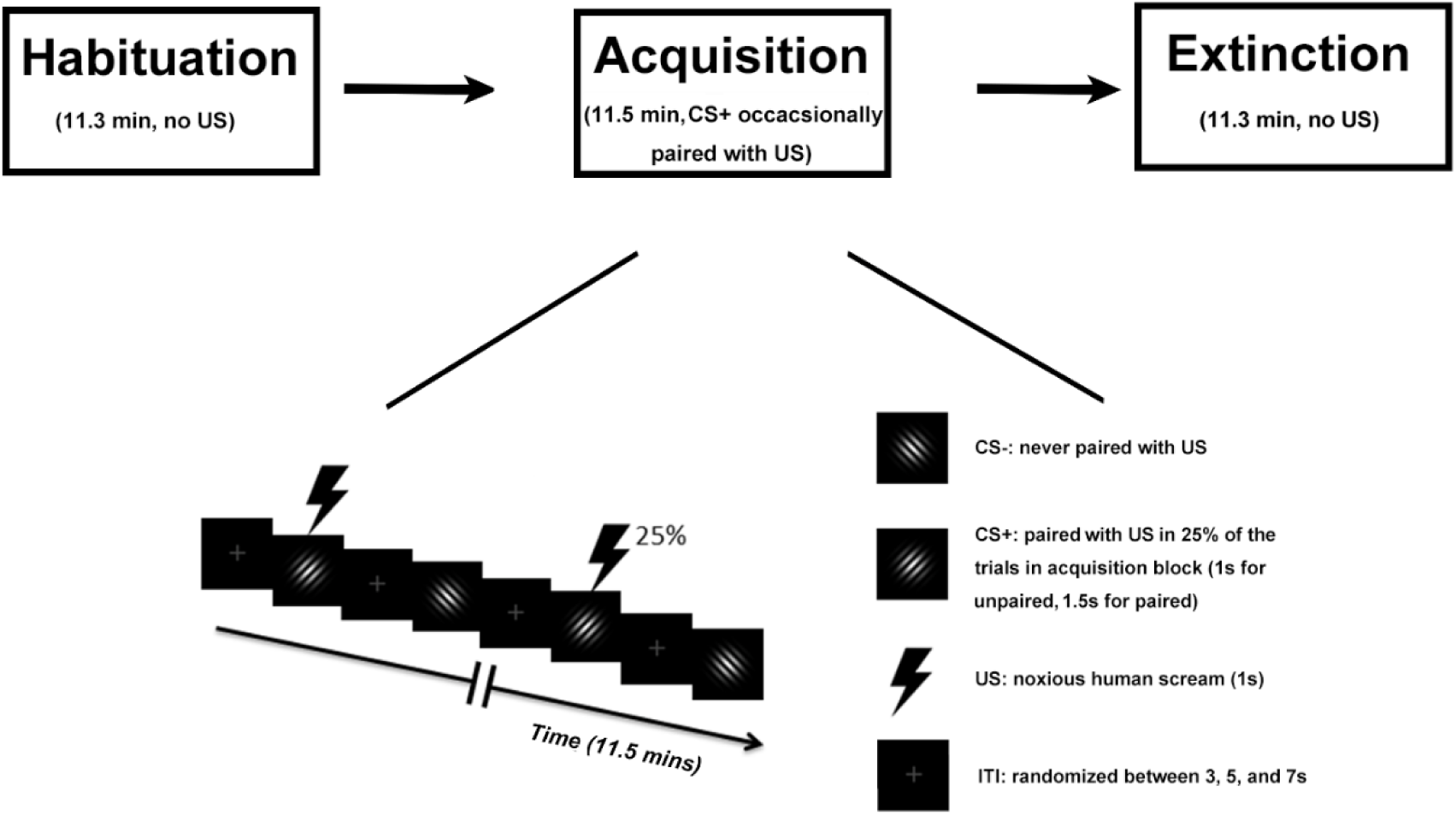
Experimental paradigm. Top: Temporal order of the three blocks. Bottom: timeline and stimuli used during the acquisition block. For the habituation block and the extinction block, stimulus types, stimulus duration, and inter-trial interval (ITI) were the same, except that no US was presented.

#### Paradigm

There were three blocks: habituation, acquisition and extinction (Figure 1). Each block comprised of 120 trials and lasted about 12 minutes. During the acquisition block, one Gabor patch was designated as CS+ and the other as CS-. In the habituation block, which preceded the acquisition block, the two Gabor patches occurred with equal probability in a pseudo-random order, determined by a procedure in which the two Gabor patches were randomly (toss of a fair coin) drawn from two pools without replacement, under the constraint that not more than 2 CSs of one kind (future CS+, future CS-) appeared in direct sequence, as is typical in fear conditioning work (Lonsdorf et al., 2017). During acquisition, the same pseudo-randomization was again applied to result in a different order with the same constraints as described for habituation. In addition, acquisition always started with a CS+ trial, and the first 4 CS+ stimuli were always paired with the US to facilitate contingency learning. Subsequently, 25% of CS+ stimuli were paired with the US. CS-stimuli were never paired with the US. For analysis, paired CS+ trials were not included due to fMRI contamination by US evoked responses. For notational simplicity, in what follows, we use the term CS+ trials when referring to unpaired CS+ trials in which no US occurred. In the extinction block, which followed the acquisition block, the stimuli and procedure were the same as the habituation block, i.e., the pseudo-randomization procedure was again applied to result in a pseudo-random order with the constraint that no more than two CSs of the same type appeared in a row. We note that, within a given block, the order of trials was the same for each subject to facilitate trial-by-trial averaging across subjects, which is essential for analyzing the temporal dynamics of conditioning across trials at a population level. For each of the three blocks, the inter-trial interval (ITI) was randomized between 3, 5, and 7 secs; see Figure 1.

### Data Acquisition

#### Functional MRI data

Functional MRI (fMRI) images were acquired on a 3-Tesla Philips Achieva whole-body MRI system (Philips Medical Systems, Netherlands) using a T2*-weighted echoplanar imaging (EPI) sequence (echo time (TE) = 30ms; repetition time (TR) =1980 ms; flip angle=80°). Each whole-brain volume consisted of 36 axial slices (field of view: 224mm; matrix size: 64×64; slice thickness: 3.50 mm; voxel size: 3.5×3.5×3.5mm). A T1-weighted high resolution structural image was also obtained from each participant. For one subject, the fMRI data during habituation were not properly saved to the disk, and the data from 17 subjects were used for fMRI analysis for the habituation block. For all other analyses, fMRI data from all 18 subjects were used.

#### EEG data

EEG data was recorded simultaneously with fMRI using a 32-channel MR-compatible EEG system (Brain Products GmbH, Germany). Thirty-one sintered Ag/AgCl electrodes were placed according to the 10–20 system with the reference channel being FCz during recording. One additional electrode was placed on the participant’s upper back to monitor the electrocardiogram (ECG). ECG data was used to enable heart rate (HR) analysis and to aid in the removal of the cardioballistic artifacts. The impedance from all scalp channels was kept below 10 kΩ during the entire recording session as recommended by the manufacturer. The online band-pass filter had cutoff frequencies at 0.1 and 250 Hz. The filtered EEG signal was then sampled at 5 kHz and digitized to 16-bit. The EEG recording system was synchronized with the scanner’s internal clock, which, along with the high sampling rate, was essential to ensure the removal of the MRI gradient artifacts.

### Regions of Interest

Three regions of interest (ROIs) were considered: the primary visual cortex or V1 ROI (Figure 2A), the right temporal parietal junction (rTPJ) ROI (Figure 2B), and the right amygdala ROI (Figure 2C). The V1 ROI was bilateral and defined using a recently published template of retinotopic regions of the visual cortex (Wang et al., 2014); this ROI contained 473 contiguous voxels. The rTPJ ROI was defined to be a 6mm sphere centered at the previously published coordinates of TPJ (Geng and Vossel, 2013); this ROI contained 33 voxels. The right amygdala ROI was chosen to be a 6mm sphere centered at the previously determined peak-activation voxel from contrasting US against CS-(Yin et al., 2018); this ROI contained 33 voxels. The left amygdala was not activated in this contrast and thus not considered further.

**Figure 2:**
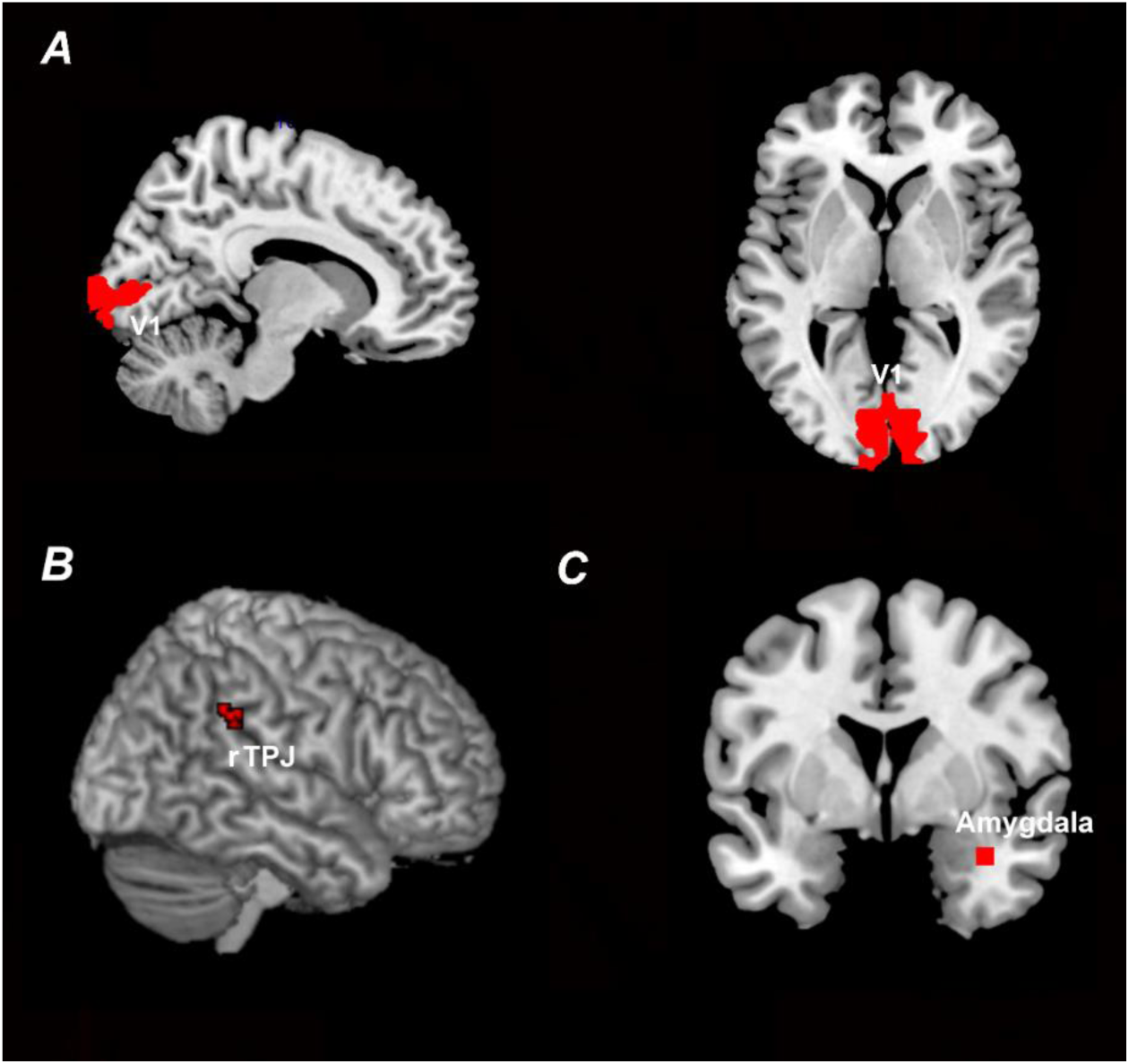
ROI definition. A) V1 ROI defined according to a recently published retinotopic atlas of the visual cortex by Wang et al. (2014). B) rTPJ ROI defined according to previously published coordinates (Geng and Vossel, 2013). (C) Right amygdala ROI defined according to previously published coordinates based on US activation (Yin et al., 2018).

### Data Processing

#### FMRI data preprocessing

All fMRI analyses were performed in SPM (http://www.fil.ion.ucl.ac.uk/spm/). Preprocessing steps included slice timing, motion correction, and normalization to the Montreal Neurological Institute (MNI) template. Normalized images were spatially-smoothed with a 7mm FWHM (Full Width at Half Maximum) Gaussian kernel. This spatial smoothing step was omitted for the representational similarity analysis (RSA) analysis to better preserve spatial patterns. Global scaling was applied to remove the global signal from the BOLD time series (Desjardins et al., 2001). The BOLD time series were then high-pass filtered with a cutoff frequency at 1/128 Hz.

#### EEG data preprocessing

There are two major sources of MRI-related artifacts in EEG recorded simultaneously with fMRI: the gradient artifacts and the cardioballistic artifacts. Gradient artifacts were removed by subtracting an average artifact template from the data set as implemented in Brain Vision Analyzer 2.0 (Brain Products GmbH, Germany). The artifact template was constructed by using a sliding-window approach which involved averaging the EEG signal across the nearest 41 consecutive volumes. The cardioballistic artifacts were also removed by an average artifact subtraction method (Allen et al., 1998). In this method, the R peaks were first detected in the ECG recordings by the algorithm in Brain Vision Analyzer, and then visually inspected to ensure accuracy. The appropriately detected R peaks were utilized to construct a delayed average artifact template over 21 consecutive heartbeat events. The cardio-ballistic artifacts were then removed by subtracting the average artifact templates from the EEG data. After these two steps, the EEG data were band-pass filtered between 0.5 and 50 Hz, down-sampled to 250 Hz, re-referenced to the average reference (Nunez et al., 1997), and exported to EEGLAB (Delorme and Makeig, 2004) for analysis.

#### Heart rate (HR) analysis

The RR intervals was estimated from the ECG data and transformed into instantaneous HR (inverse of RR interval). The time range from 1-s prestimulus to 5-s poststimulus was divided into 1s bins, and each instantaneous HR was weighted proportionally to the fraction of the bin it occupied (Gatchel and Lang, 1973; Graham, 1980) to yield stimulus-locked HR change times series within a trial. This single-trial stimulus-locked HR change time series was then averaged across all trials within a block to assess how, on average, CS+ and CS-affected stimulus-locked HR changes in habituation, acquisition and extinction.

In addition, for each of the three blocks, the time courses of relative HR changes over trials were obtained by taking the stimulus-locked HR change in the interval (0.5 s, 1.5 s) from each trial, smoothing it over trials for CS+ and CS-separately using a Gaussian kernel (bandwidth=12) to yield CS+ HR curve and CS-HR curve, and computing the difference by subtracting the CS+ HR curve from the CS-HR curve.

#### Single-trial estimation of BOLD response

The BOLD response was estimated on a trial-by-trial basis using the beta series method (Rissman et al., 2004; Rissman et al., 2008). Details can be found in our previous paper (Yin et al., 2018). Briefly, in this method, every stimulus was associated with a separate regressor in the general linear model (GLM). Rigid body movements were included as regressors of no interest. Solving the GLM yielded a beta value for each trial in each voxel.

#### Representational similarity analysis (RSA)

Voxel representations of CS+ and CS-can be studied using RSA, a MVPA method (Visser et al., 2011; Visser et al., 2013). To maximally retain information at a finer spatial scale (Dunsmoor et al., 2014), we applied the beta series method to the BOLD time series prior to spatial smoothing to obtain single trial beta values. For a given ROI, a vector was created from the beta values of all the voxels to represent the spatial pattern in response to a single presentation of a stimulus; the length of the vector equaled the number of voxels in that ROI. Reference representations of CS+ and CS-for the V1 ROI were generated from averaging the single trial multivoxel patterns across all the trials (60 each) in the habituation block. During acquisition and extinction, to generate the time course of neural representational changes over trials (i.e., similarity curve), a sliding window approach was adopted, in which the time window used was 5 trials in duration and the step size was 1 trial. After the moving average (5-trial average), each trial-averaged vector in acquisition and extinction was correlated with its reference representation derived from habituation to assess pattern similarity. The correlation coefficients were Fisher-z transformed, averaged across participants, re-transformed back to correlation coefficients, and plotted as a function of time-on-task to yield the time course of changes in neural representations of CS+ and CS-. The slope of the time course was estimated by linear fit for each individual subject’s similarity curve, and taken as the rate of change in neural representations for that subject. A paired t-test was used to compare the slopes between CS+ and CS-across participants.

#### Pattern sparsity analysis

To investigate the changes in stimulus-evoked BOLD patterns vis-à-vis the changes in stimulus-evoked BOLD magnitude during acquisition, we quantified the change in sparsity of the voxels in the representational pattern evoked by CS+ and CS-. First, to assess the broad temporal change across acquisition, we divided acquisition into an early time period (t<5.6 mins) and a late time period (t>5.6 mins). Second, for the stimulus type (CS+ or CS-) showing significant pattern change relative to habituation, we counted the voxels that represented this stimulus type (i.e., representational voxels) for each time period. Voxels entered this count only if they met all of the following 3 requirements: (1) It showed larger averaged activity for this stimulus type (e.g., CS+) than the other type (e.g., CS-), across trials; (2) The mean of the betas (across CS+ or CS-trials) from this voxel must be greater than the mean of the betas from all the voxels within that ROI; and (3) The standard deviation of the betas from this voxel across trials must be less than the mean of the standard deviation of the betas from all the voxels within that ROI. Thus, a representational voxel defined this way was a voxel that was consistently and selectively enhanced for a given stimulus type across trials, and across neighboring voxels. Finally, the number of representational voxels and the averaged betas within these voxels were compared between early and late period of acquisition to assess the changes in stimulus-evoked representational patterns and in stimulus-evoked BOLD magnitude. The same analysis was also applied to the extinction block to examine whether the sparsified neural representations of conditioned threat persisted over the extinction phase of the experiment.

#### EEG alpha ERD estimation

Event-related desynchronization (ERD) of posterior alpha oscillations (8 to 12 Hz) was taken as an indicator of visual activation and attention orienting. Alpha ERD was estimated for each trial as follows. First, the EEG signal was epoched from -1s to 2s with 0s denoting the onset of the stimulus. Second, the EEG signal within each epoch was divided into overlapped moving windows with 200ms in duration and stepped forward in 20ms increment. Third, the EEG data in each window was zero–padded to 5 times its original length (250 points after padding) to enhance spectral resolution from 5Hz to 1Hz. Fourth, the EEG power spectrum for each window was calculated using a nonparametric multi-taper approach with 3 tapers (Mitra and Pesaran, 1999), and the alpha power was estimated by averaging the power spectrum between 8 and 12 Hz. The baseline was calculated as the alpha power at stimulus onset across all trials. The single trial alpha ERD was calculated by subtracting baseline alpha and dividing the difference by baseline alpha. For a given post-stimulus time window, alpha ERD could be plotted as a function of acquisition trials, and the slope of this function obtained from a linear regression analysis provided a rate of change of visual attention engagement. A paired t-test was used to compare the slopes between CS+ and CS-category across participants. It is worth noting that the present experimental paradigm lacks an attention manipulation. Using alpha ERD as an index of visual attention engagement is indirect and relies on prior research (Klimesch et al., 2011; Auksztulewicz et al., 2017).

#### Alpha-BOLD correlation

To assess which regions of the brain (rTPJ versus right amygdala) modulated alpha power, two analyses were carried out for the acquisition block: across-participant correlation analysis and across-trial correlation analysis. Across-participant correlation was computed as the correlation coefficient between differential occipital alpha ERD (CS+ minus CS-) averaged across trials within the acquisition block and differential beta values from rTPJ or right amygdala (CS+ minus CS-) averaged across trials within the acquisition block. Across-trial correlation was assessed by correlating the single trial alpha ERD averaged across subjects and single trial BOLD beta values averaged across subjects. We sought converging evidence between these two types of analyses.

## Results

### Heart rate changes

As shown in Figure 3A and 3B, in both habituation and extinction, the average event-related HR changes did not differ between CS+ and CS-. During acquisition, greater HR deceleration was observed following CS+ compared to CS–, demonstrating that participants acquired the contingencies of the experiment, and that they exhibited defensive orienting to the CS+. Figure 3C shows the time course of relative event-related HR change (CS+ minus CS-) over trials for the three blocks. During habituation, as expected, there was no systematic trend in differential HR time course across trials. For acquisition, greater CS+-related HR deceleration was apparent in the early part of the block, and the difference gradually diminished as learning progressed, and disappeared toward the end of the block (Yin et al., 2018). There was no systematic trend in event-related HR change between CS+ and CS-over the entire extinction block.

**Figure 3:**
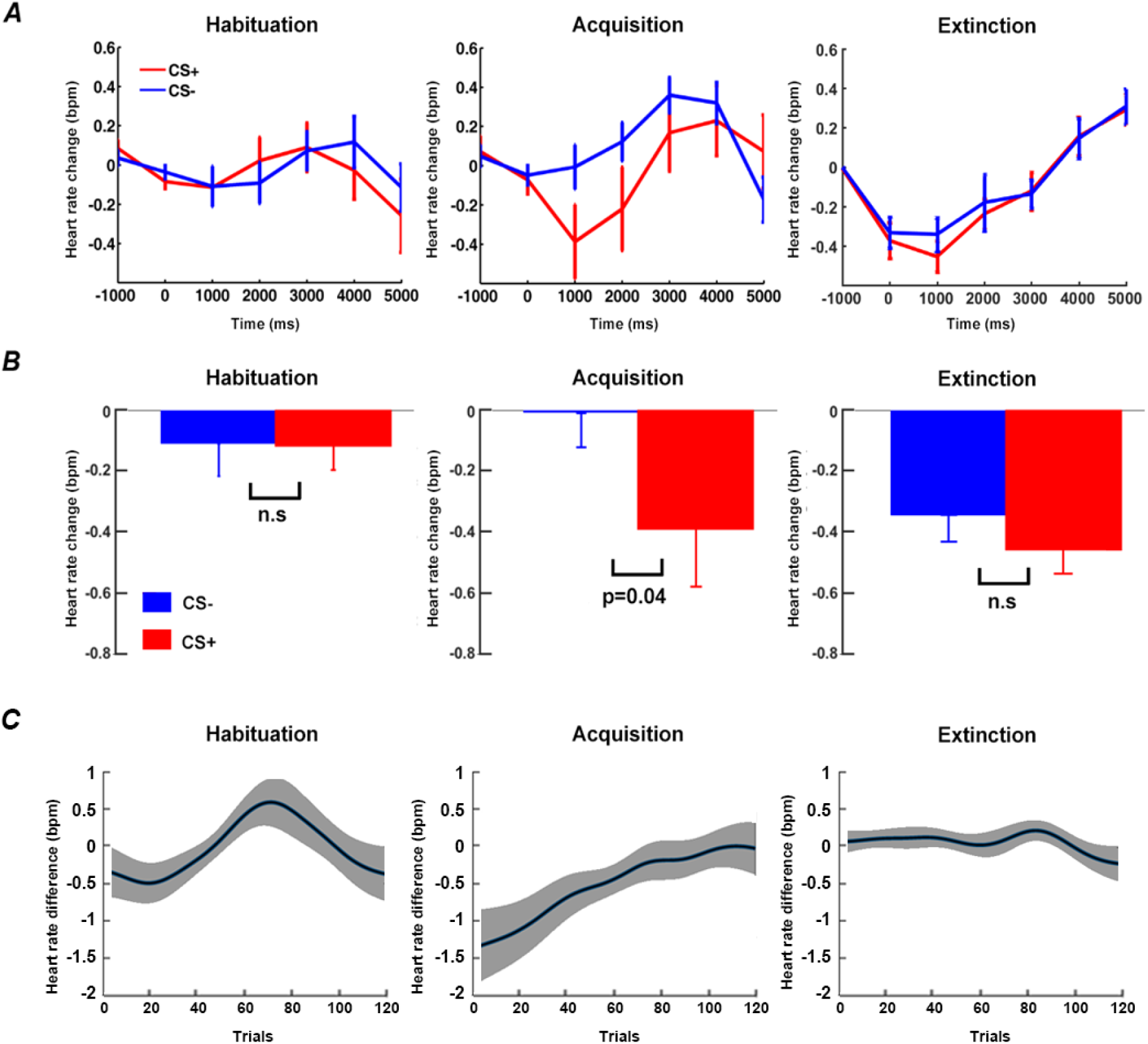
Heart rate (HR) analysis. A) Event-related HR changes during habituation, acquisition and extinction. B) Statistical comparison of HR between CS+ and CS-at time=1 s (0.5 s to 1.5 s). C) Time course of relative event-related HR changes (CS+ minus CS-) over trials in habituation, acquisition and extinction. Note: Figure 3C (left) and 3C (middle) are adapted from Yin et al., (2018) and included here for comparison with Figure 3C (right).

### Dynamic changes of neural representations of CS+ and CS-in acquisition

Reference representations for CS+ and CS-in V1 were obtained by averaging single trial BOLD responses to CS+ and CS-across habituation trials. During acquisition, CS+ and CS-evoked patterns in V1 were correlated with their respective reference representational patterns using a moving window approach, to yield the time courses of RSA pattern similarity changes (similarity curves); see Figures 4A and 4B for example similarity curves from an individual subject. Across participants, the RSA similarity curve for CS+ showed a decreasing trend, while the similarity curve for CS-varied unsystematically, resulting in a flat average slope (Figures 4C and 4D). This demonstrates that the patterns evoked by CS+ during acquisition were systematically departing from its reference representation pattern, whereas the CS-evoked patterns did not exhibit any systematic change. The rate of pattern similarity change for each individual, indexed by the slope of the linear fit to the similarity curve, is shown in Figure 4C. Across participants, as shown in Figure 4D, the slopes of CS+ RSA similarity curves were significantly different from the slopes of CS-RSA similarity curves (p<0.01).

**Figure 4:**
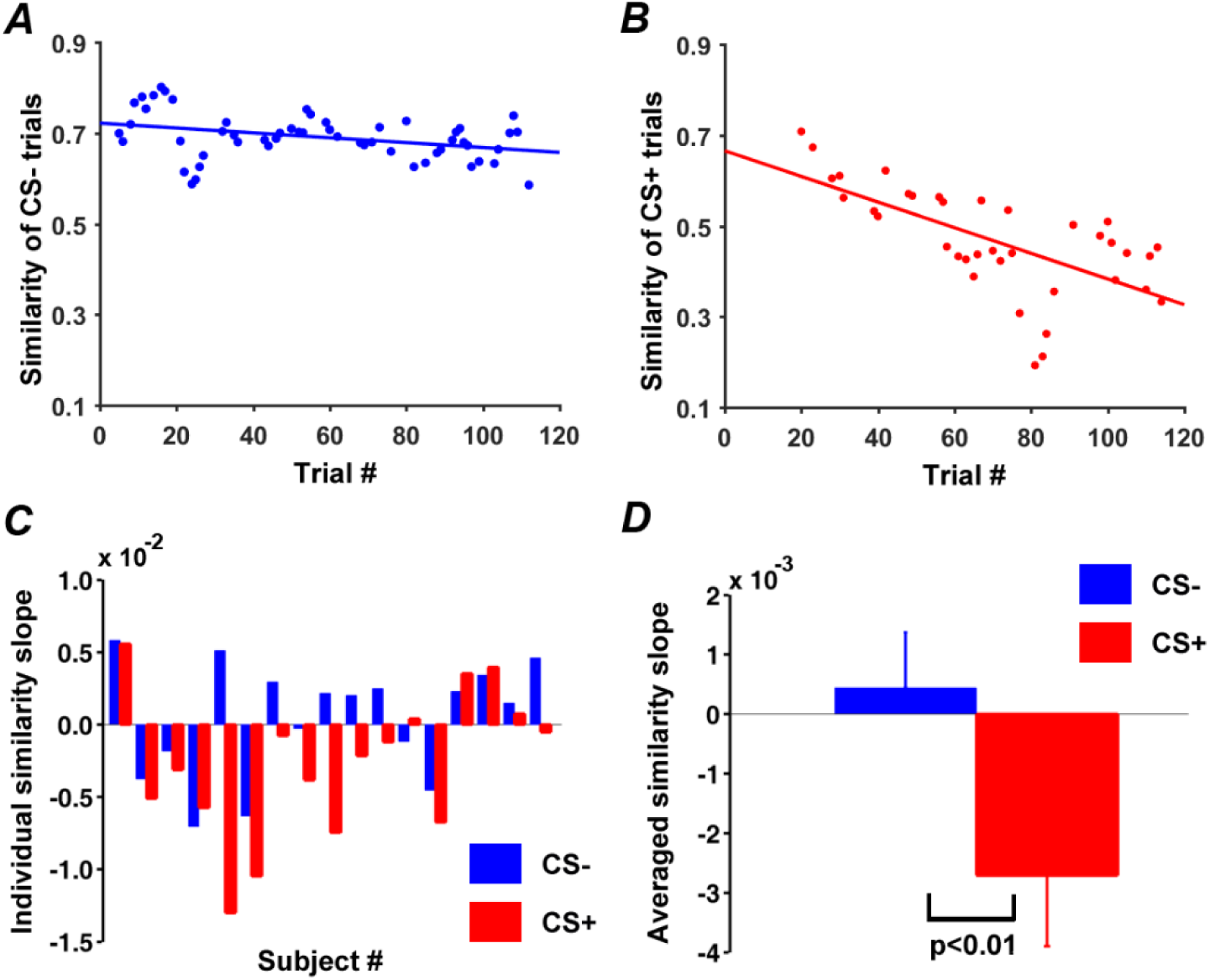
Pattern similarity changes during acquisition in V1. A) Time course of pattern similarity change in V1 for CS-trials (Subject 8 in C). B) Time course of pattern similarity change in V1 for CS+ trials from the same subject. C) Slopes of linear fits to pattern similarity curves for each participant. D) Slopes of similarity curves between CS+ and CS-in V1 were significantly different.

### Changes in pattern sparsity during acquisition

To more closely examine the acquisition-related CS+ pattern changes over time, we divided the habituation block and the acquisition block into an early period and a late period. For each time period, the V1 representational voxels (defined in Methods: pattern sparsity analysis) for CS+ were counted and shown in Figure 5A for habituation and Figure 5C for acquisition, and the averaged betas within these voxels, representing BOLD response magnitude, were calculated and plotted in Figure 5B for habituation and Figure 5D for acquisition. For habituation, the number of representational voxels was not significantly different between the early and the late period (Figure 5A), whereas the average BOLD response magnitude became significantly lower in the late period (p=0.01) (Figure 5B). For acquisition, the number of representational voxels was lower in the late period relative to the early period (p<0.004) (Figure 5C), while the BOLD response magnitude did not undergo significant change (Figure 5D). Figure 5E illustrates the multivoxel patterns evoked by CS+ with the color of each cube (i.e., voxel) reflecting the beta value of that voxel; in the late period of acquisition, CS+ was represented by fewer voxels compared to the early period of acquisition.

**Figure 5:**
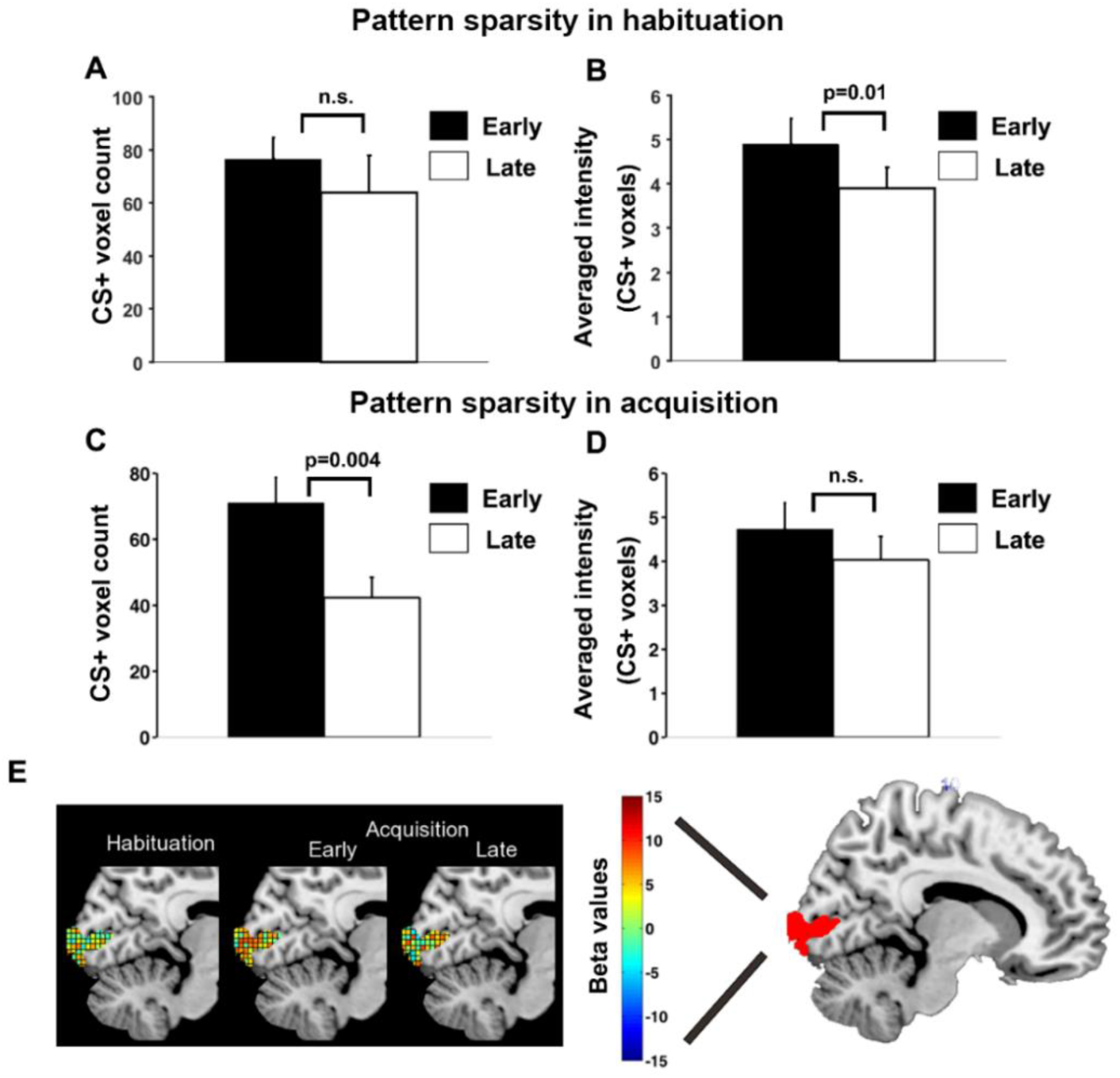
Pattern sparsity analysis for CS+ trials. A) No significant difference in number of representational voxels for CS+ in V1 between early and late habituation. B) BOLD activity magnitude was significantly lower in late habituation than early habituation. C) Number of representational voxels for CS+ in V1 was significantly lower in late acquisition than early acquisition. D) No significant difference in BOLD activity magnitude between early and late acquisition. E) Schematic illustration of increasing sparsity observed during CS+ trials over time: CS+ evoked multivoxel patterns of beta values in habituation, early acquisition and late acquisition.

### EEG alpha-band activity

Event-related alpha-band power (8 to 12 Hz) was shown in Figure 6A. Quantifying alpha ERD using average alpha power in the interval 600 to 1000 ms, there was no significant difference in alpha power between CS+ and CS-in habituation or in early period of acquisition, but alpha power was significantly lower following CS+ relative to CS-in late period of acquisition (t(17)=-17.73, p<0.01) (Figure 6B). In line with this finding, a paired t-test showed greater differential (CS+ minus CS-) alpha ERD in the late compared to the early period of acquisition (t(17)=2.75, p=0.01); see Figure 6B. To further quantify these cross-trial dynamics, we computed the time course of alpha ERD changes using the moving window approach mentioned earlier, and estimated the slope of the linear fit to alpha ERD changes across CS+ trials and CS-trials. A Wilcoxon signed-rank test indicated that the resulting slopes differed significantly (Z=2.50, p=0.01); see Figure 6C. CS+ trials showed more alpha power decrease over time than CS-trials (p<0.01).

**Figure 6:**
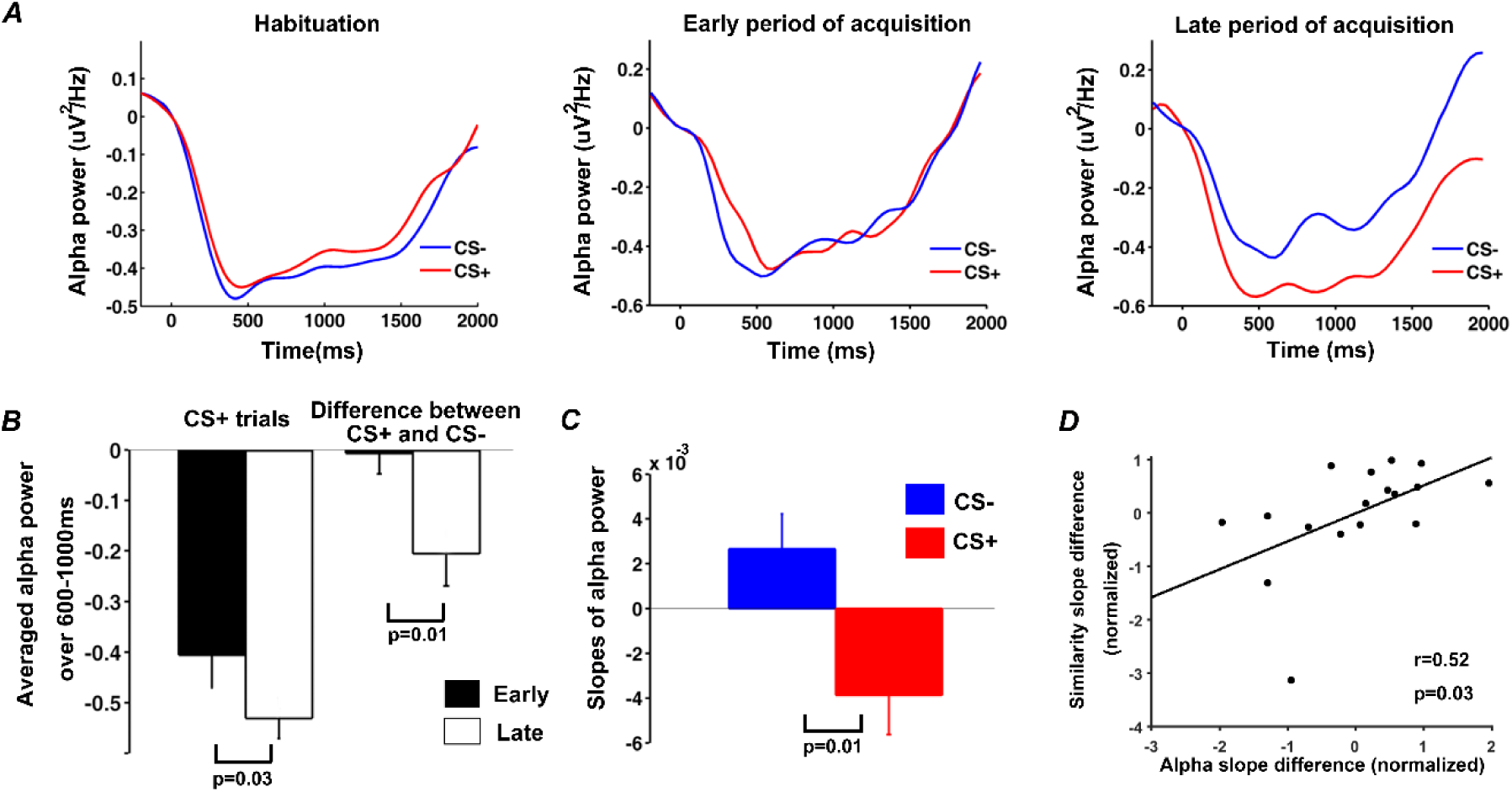
Event-related alpha desynchronization during habituation and acquisition. A) Alpha-band (8-12 Hz) power averaged across CS+ and CS-trials during habituation, the early period of acquisition, and the late period of acquisition. B) CS+-evoked alpha ERD and the difference in CS+ and CS-alpha-band power between early and late acquisition periods. C) The slope of linear fit to the time course of alpha-band power across acquisition trials. D) Relation between the rate of event-related alpha-band power decrease and the rate of pattern similarity change in V1 (each point in the plot represents one participant).

### Alpha ERD change and BOLD pattern similarity change during acquisition

Exploring the relationship between changes in alpha ERD and changes in CS+ evoked patterns in V1 during acquisition, we observed a positive correlation at r=0.52 and p=0.03, between the differential slope of alpha power event-related desynchronization and the differential slope of pattern similarity curve; see Figure 6D. This finding suggests that as aversive learning progressed, participants with more pronounced representational voxel pattern changes in V1 tended to show progressively stronger alpha ERD.

### Alpha-BOLD correlation during acquisition

Concurrent recordings of EEG and fMRI afforded the opportunity to examine the sources of modulatory signals for alpha ERD. In acquisition, as shown in Figure 7, alpha power desynchronization was found to be significantly negatively correlated with the BOLD from rTPJ both across participants (r=-0.51, p=0.03; Figure 7A) and across trials (r=-0.22, p=0.02; Figure 7B). However, no correlation was found between alpha power desynchronization and the BOLD in right amygdala both across participants and across trials (Figures 7A and 7B).

**Figure 7:**
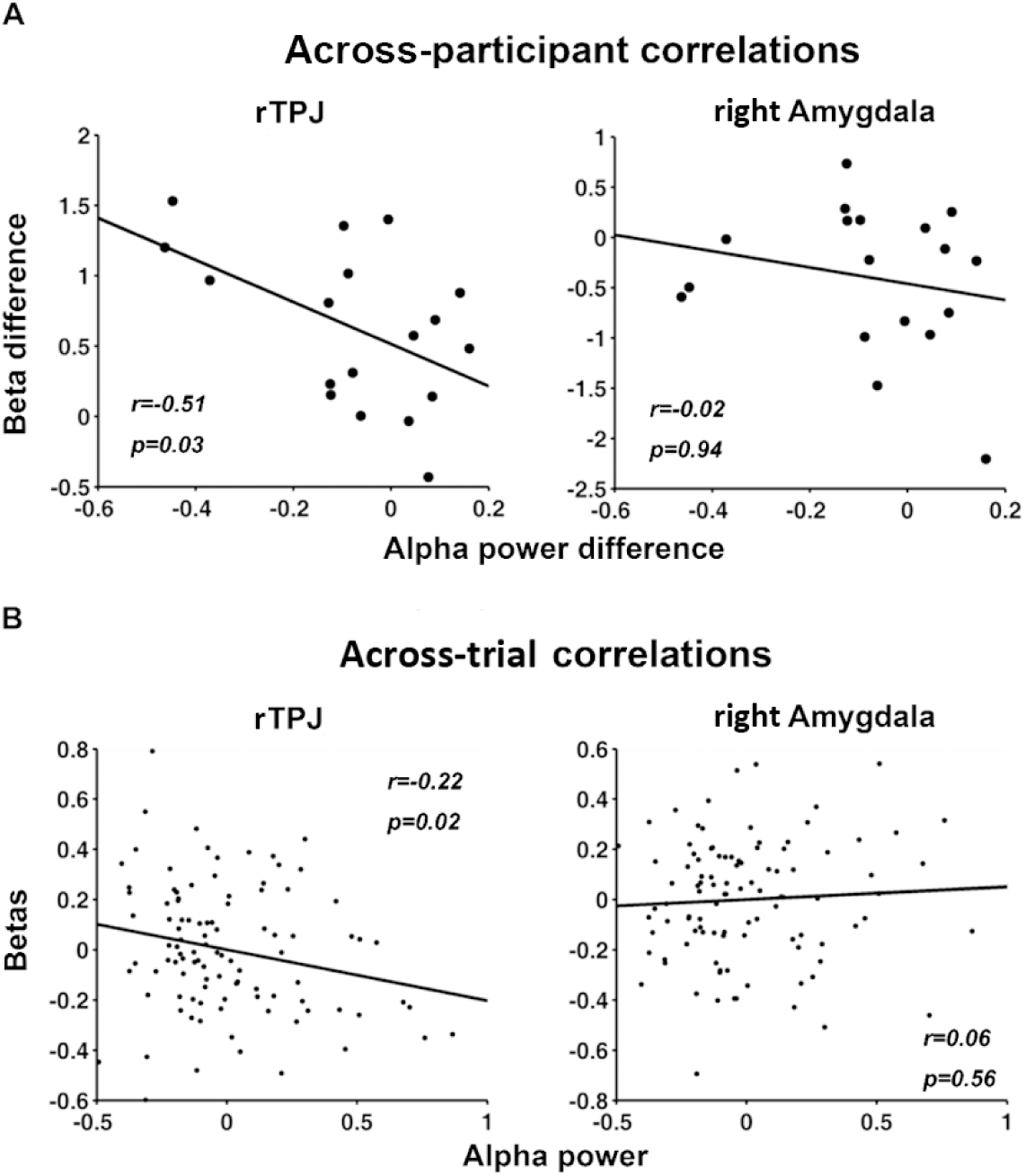
EEG-BOLD coupling in acquisition. A) Across-participant correlations between alpha-band ERD and BOLD in the rTPJ, a core structure of the ventral attention network, and the right amygdala. A negative correlation was observed between alpha ERD difference (CS+ minus CS-) and the difference in rTPJ beta values (CS+ minus CS-). No correlation was observed between alpha ERD difference and the estimated beta difference in right amygdala. Each point in the plots represents a participant. B) Across-trial correlations between alpha ERD and BOLD in rTPJ and right amygdala. Similar to A), there was a significant negative correlation between the single trial alpha power and the single trial betas from rTPJ, and no correlation between the single trial alpha power and the single trial betas from right amygdala. Each point in the plots represents a trial.

### Neural dynamics during extinction

We carried out similar analyses for the extinction data. During extinction, as shown in Figure 8, slopes of pattern similarity change time course were not different between CS+ and CS-; the number of representational voxels for CS+ and CS+ evoked BOLD intensity were not different between early and late periods; and alpha ERD for CS+ and CS-were not significantly different between early and late extinction. A closer inspection of Figures 5C and 8C revealed that, the number of representational voxels for CS+ during early extinction (47.5±4.5) and the number of representational voxels for CS+ in late acquisition (42.5±5.5) were not significantly different (p>0.8), suggesting that the sparser neural representations of CS+ reached at the end of acquisition persisted in extinction.

**Figure 8:**
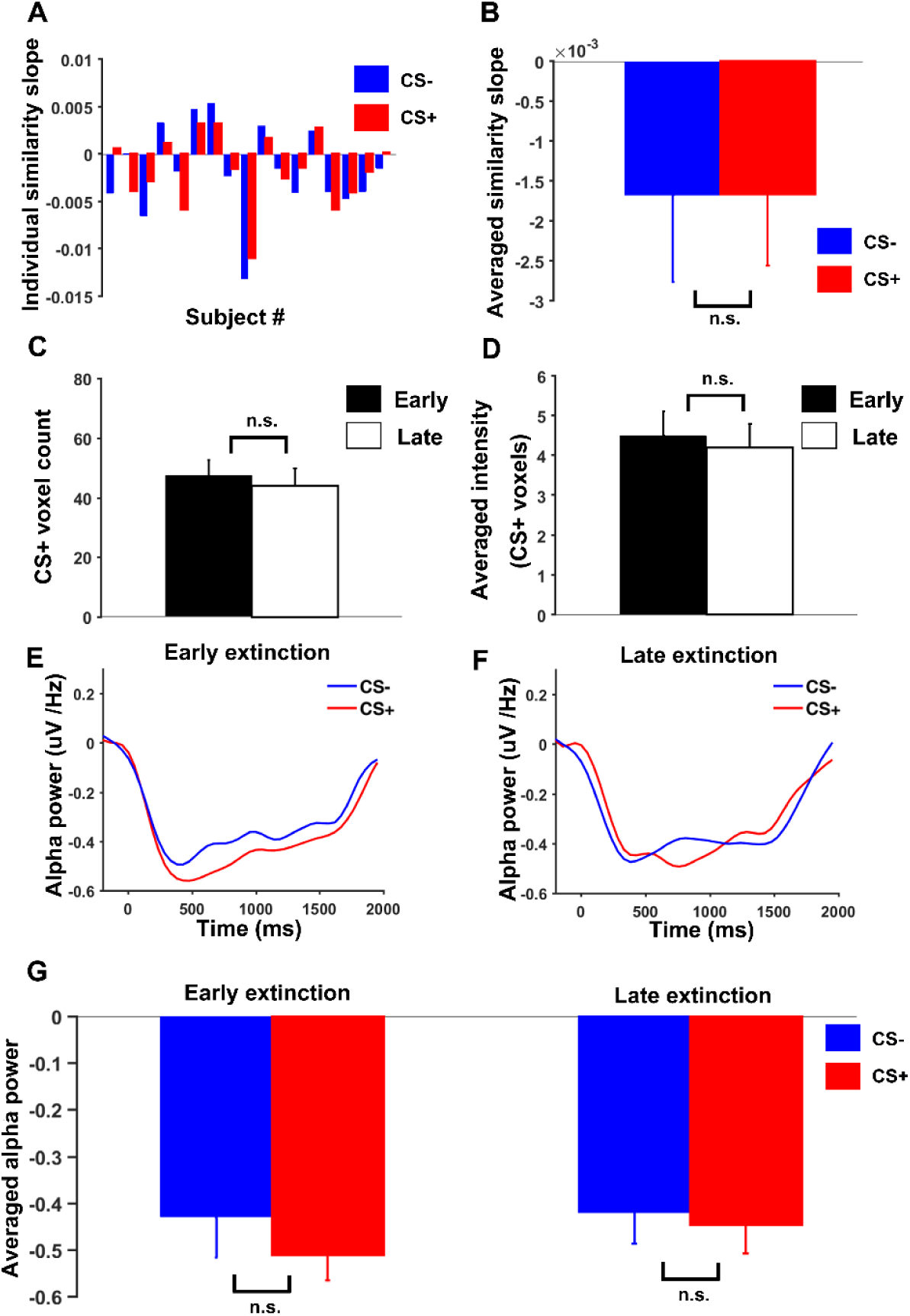
Neural dynamics in V1 during extinction. A) Slopes of linear fits to pattern similarity curves for each participant. B) Slopes of similarity curves were not significantly different between CS+ and CS-. C) No significant difference in number of representational voxels for CS+ between early and late extinction. D) No significant change in BOLD activity magnitude for CS+ between early and late extinction. Alpha-band ERD for CS+ and CS-trials during E) early extinction (p=0.15) and F) late extinction (p=0.29). No significant difference in alpha ERD between CS+ and CS-in either G) early or H) late extinction.

## Discussion

In classical fear conditioning, a neutral stimulus (CS+), through association with an aversive stimulus, comes to elicit defensive responses in the absence of the original aversive stimulus. The sensory response to CS+ also undergoes systematic changes in this process. Here, we found that: 1) in primary visual cortex (V1), the representational voxel patterns evoked by the CS+ became sparser, and this sparsification persisted over extinction; 2) alpha ERD became more pronounced as learning progressed, suggesting heightened engagement of attention and arousal systems; 3) the rate of change in V1 representation of CS+ was positively related to the rate of change in alpha ERD; and 4) EEG alpha band activity was coupled to BOLD activity in rTPJ but not to BOLD activity in right amygdala.

### Sharpened Visual Representation

Electrophysiological studies in humans have found visuocortical amplification of conditioned threat cues (Moratti et al., 2006; Stolarova et al., 2006; Miskovic and Keil, 2012; Thigpen et al., 2017), accompanied by heightened inter-trial and inter-site phase locking over primary visual cortex (Keil et al., 2007, McTeague et al., 2015). The present study suggests that such changes reflect a sparsification process, in which visual features associated with recurring, predictable threat are increasingly represented by sharpened, efficient, and internally tightly coupled visuocortical networks, rather than by a generally heightened visual population response. Specifically, we found that fewer voxels contributed to the representation of CS+ as learning progressed, whereas the BOLD magnitude evoked by CS+ did not. Sparsification of voxel patterns is conceptually consistent with notions of sharpened, efficient representations emerging as a function of Hebbian associative mechanisms. In electrophysiological work, such networks would be expected to show heightened temporal accuracy and phase stability across trials, which is what has been observed in previous studies (Miskovic and Keil, 2012). Notably, sparsification persisted throughout extinction, despite evidence showing that the selective HR orienting response to the CS+ was extinguished. The observation that changes in visuocortical activity are more resistant to extinction than autonomic or behavioral indices is consistent with studies on experimental animals (Headley and Weinberger, 2011; Moran and Katz, 2014) as well as human participants (McTeague et al., 2015). These studies have shown sustained sensory learning and sensory plasticity during extinction, instead of returning to a pre-conditioning, naïve, state (for a review, see McGann, 2015). Sparsification has been discussed as a key aspect of such ongoing plasticity, because it minimizes metabolic cost while enabling specific and efficient representations of predictable threat cues (Miskovic and Keil, 2012; McGann, 2015).

Alternatively, a body of research has suggested that repeating visual stimuli produces neural activity reduction in the visual cortex, called repetition suppression, potentially accompanied by sharpened representations (e.g., Gruber and Müller, 2002; Gruber et al., 2004). A repetition suppression effect alone is however unlikely to explain the present set of findings, because 1) decreasing activity induced by viewing the same stimulus repeatedly has been primarily observed with familiar, meaningful objects and scenes, whereas repetition of unfamiliar stimuli devoid of rich semantics (such as the gratings used in the present study) may lead to an increase of neural activity under this perspective (Conrad et al. 2007), 2) the number of repetitions of the CS+ and CS-was equal across the trial types (60 trials each), thereby ruling out the effect of uneven stimulus exposure, and 3) the voxel pattern difference prompted by the two stimuli persisted during extinction training in which both CS+ and CS- were shown in identical fashion, with no US given. Furthermore, whereas the number of voxels that selectively represented the CS+ became smaller as learning progressed, the overall BOLD activity within these voxels did not change. One may argue that the change and sparsification of the V1 BOLD patterns may simply be a reflection of diminished engagement with the threat cue over the course of the experimental session. The increased alpha ERD for the CS+ with learning, however, is more consistent with the notion that attention is increasingly directed to the threat cue, contradicting a selective disengagement hypothesis. Together, the present data support the hypothesis that associative learning selectively shapes visuocortical representations of threat in a way that promotes sparse, sharpened coding of the critical stimulus features (Kok et al., 2012; Ibrahim et al., 2016).

### Defensive Orienting and EEG Alpha Power Reduction

Previous work has shown that the cardiac orienting response to threat, measured as phasic heart rate deceleration when viewing the CS+, is attenuated as learning progresses (Sokolov, 1963; Bradley, 2009). This adaptation in HR orienting is concomitant with adaptation in canonical fear circuits, including the amygdaloid complex (Yin et al., 2018). By contrast, in the present study, EEG alpha desynchronization— associated with visual activation and attentive stimulus processing—showed an increase (sensitization) during the course of acquisition. This is consistent with the long-held notion that behavioral, autonomic, and neurophysiological responses to threat are not linearly related (Lang, 1979), reflective of their different adaptive functions in addressing the threat.

A large body of research has shown that the extent of event-related alpha power reduction over visual areas co-varies with the motivational significance (task-relevance) and/or perceptual saliency of the event (Ruby et al., 2013). Thus, the present finding that alpha ERD becomes stronger with conditioning suggests that the selective/attentive processing of the CS+ is increasing, not decreasing as learning progresses. Supporting this interpretation, a previous study found the adaptation of limbic brain areas to be accompanied by increased engagement of visual cortex during fear conditioning (Lithari et al., 2016). Such persistent visuocortical engagement with the threat cue may be particularly adaptive in conditioning regimes with intermittent pairing, in which not all CS+ trials include a US presentation, promoting exploration behavior and scanning of the environment for contingency cues—a speculation that is readily testable in future research.

### Sources of Visuocortical Changes during Fear Conditioning

Most contemporary viewpoints agree that heightened visuocortical responses result from interactions between visual and extra-visual brain regions, with the latter conveying modulatory signals that selectively heighten the gain of visual neurons, individually or at the population level. Two candidate circuits for providing re-entrant modulatory feedback to visual cortex have received the most attention in the literature: the amygdala and the ventral attention network. Two mechanisms have been proposed for amygdalofugal modulations of the visual system. One is through its projections to earlier levels of the visual pathway including primary and secondary visual cortices to enhance perceptual processing of emotional stimuli (Amaral et al., 2003). The other is through its connections with higher order attentional modulation areas such as the intraparietal sulcus (Armony and Dolan, 2002) and vlPFC (Ghashghaei et al., 2007). Consistent with earlier work testing the amygdalofugal re-entry hypothesis in fear conditioning (Petro et al., 2017), the present study did not find support for the notion that hemodynamic activity in the amygdaloid complex co-varies with selective visuocortical processing of the CS+, neither at the level of BOLD nor at the level of scalp-recorded electrophysiology. Targeted studies in the animal model are needed to characterize the role of the amygdala in biasing visuocortical processes during fear conditioning.

The rTPJ is a key node of the ventral attention network which has been proposed to mediate the allocation of attention in response to the presence of salient sensory stimuli (Fox et al., 2006; Vossel et al., 2014). For example, BOLD activity in areas within the VAN such as the rTPJ is modulated by tasks that require participants to selectively attend to events varying in hedonic valence and/or arousal (Fichtenholtz et al., 2004; Lee and Siegel., 2012). Although not suitable for establishing causality, the present findings support the hypothesis (Petro et al., 2017) that, even in the absence of a cognitive task, biasing signals originating in attention-related brain regions such as rTPJ facilitate the selective visuocortical processing of conditioned threat cues. Further illustrating a dissociation of limbic and attention networks, competing macroscopic networks may be active during different phases of classical fear conditioning, with limbic and prefrontal networks being anti-correlated (Marstaller et al., 2016). Future work may address the extent to which rTPJ engagement in fear acquisition is driven by input from threat-modulated regions such as the amygdala or insula.

### Summary and conclusions

The present study showed that extensive fear conditioning prompts the emergence of sharpened, sparser pattern representations of the condition threat in visual cortex. These pattern changes were characterized by decreasing numbers of voxels showing CS+ specificity. The rate of CS+ representational pattern changes co-varied with the rate of increased CS+ evoked alpha ERD, with alpha ERD being associated with activity of rTPJ, rather than the amygdala. The sparsification of voxel patterns persisted during extinction training, in line with electrophysiological work showing lasting changes in afferent visuocortical processing after extensive fear conditioning (Thigpen, Bartsch, and Keil, 2017), despite the fact that autonomic responses to CS+ and CS-showed no difference. Together, these observations support the notion that sustained fear learning prompts plastic changes at the lowest level of visuocortical processing stream to cope with the demands posed by an ever-changing environment, and to facilitate the detection and identification of threats or opportunities.

## Acknowledgements

This work was supported by NIH grants R01MH097320 and R01 MH112558.

## Note

Conflict of interest:None declared.

